# Conversion of Diffusely Abnormal White Matter to Focal Lesions is Linked to Progression in Secondary Progressive Multiple Sclerosis

**DOI:** 10.1101/832345

**Authors:** Mahsa Dadar, Sridar Narayanan, Douglas L. Arnod, D Louis Collins, Josefina Maranzano

## Abstract

**Objectives:** 1) To automatically segment focal white matter lesions (FWML) and Diffusely abnormal white matter (DAWM), i.e. regions of diffuse abnormality observed on conventional (T2-weighted) MRI and characterize their longitudinal volumetric and normalized T1-weighted (T1w) intensity evolution, 2) To assess associations of FWML and DAWM with Expanded Disability Status Scale (EDSS) and confirmed disability progression (CDP).

**Methods:** Data includes 3951 timepoints of 589 SPMS participants followed for three years. FWML and DAWM were automatically segmented using a 2-weighted-intensity thresholding technique. Screening DAWM volumes that transformed into FWML at the last visit (DAWM-to-FWML) and normalized T1w intensities (as a marker of severity of damage) in those voxels were calculated.

**Results:** FWML volume significantly increased and DAWM volume significantly decreased as disease duration increased (p<0.001). Global EDSS scores were positively associated with FWML volumes (p=0.002), but not with DAWM volumes. Median volume of DAWM-to-FWML was significantly higher in patients who progressed (2.75 vs 1.70 cc; p<0.0001), and represented 14% of the total DAWM volume at screening, compared to 10% in patients who did not progress (p=0.001). Normalized T1w intensity values of DAWM-to-FWML were negatively associated with CDP status (p<0.00001).

**Conclusion:** DAWM transformed into FWML over time, and this transformation was significantly associated with clinical progression. DAWM voxels that transformed had greater normalized T1w intensity decrease over time, in keeping with relatively greater tissue damage evolution. Evaluation of DAWM in progressive MS provides a useful measure to evaluate therapies that aim to protect this at-risk tissue with the potential to slow progression.

## 1. INTRODUCTION

Multiple Sclerosis (MS) is an inflammatory and neurodegenerative demyelinating disease characterized by lesions that affect the white matter (WM) and gray matter (GM) of the central nervous system (CNS)^1^. The inflammatory component of the disease is more prominent in the early phases and is characterized by disruption of the blood-brain-barrier (BBB) and lymphocytic infiltration that leads to myelin and axonal damage in a focal area traditionally described as the MS plaque^1^. In most patients (approximately 80%), the inflammatory episodes subside and these focal white matter lesions (FWML) repair either partially or completely^1^. This early phase of MS is called relapsing remitting MS (RRMS), clinically translating into successive episodes of symptoms that spontaneously regress with varying degrees. As the disease evolves, the accumulation of these episodes and their incomplete recovery lead to an accumulation of lesions and progressive loss of axons, which is considered the neurodegenerative hallmark of MS, more prominent in the later stages^2, 3^. This later phase clinically translates into an accumulation of symptoms without, or un-related to, relapses and it is known as secondary progressive MS (SPMS)^4^.

During the last three decades, RRMS has been studied in depth, and its inflammatory component, in the form of FWML, has been characterized in detail, using MRI and histopathology data^3,5^. On MRI, focal lesions are relatively straightforward to detect and quantify; they appear as discrete areas of high signal intensity on T2 weighted (T2w) and fluid attenuated inversion recovery (FLAIR) images, and low signal intensity on T1 weighted (T1w) MRI contrasts^5^. Currently, there are multiple therapies that target and successfully control the development of inflammatory FWML in MS^6^.

Unfortunately, the pathophysiological mechanisms of SPMS are more elusive, and different types of lesions have been proposed as candidates that associate more significantly with progression and disability: cortical lesions (CLs)^7^, brain atrophy^8^, and diffusely abnormal WM (DAWM)^7, 9, 10^. On T2w MRI, the latter consists of mildly hyperintense areas with poorly defined edges of an intermediate value between that of normal-appearing WM (NAWM) and FWML. In histopathology, DAWM and FWML show differences in the degree of demyelination, axonal loss, and immune cell density, which are all more severe in FWML^11^.

Previous studies have shown a high correlation between normalized T1w intensity values and magnetization transfer ratio (MTR) values, which in turn have been shown to be sensitive to myelin content^12^. Hence, T1w-normalized intensities can be used as a proxy of MTR to roughly reflect the degree of demyelination of a given area: the lower the intensity value, the greater the myelin loss.

Our study takes advantage of a previously validated automated method to segment FWML and DAWM^12^ on scans of SPMS participants who were part of a three-year longitudinal study^13^. We describe the evolution of the volumes and the T1w-normalized intensity values of these two regions and explore their relationships with brain atrophy and progression of symptoms.

## 2. METHODS

### 2.1 Study Design

Our study is a retrospective, longitudinal, observational analysis to determine the volume and T1w-normalized intensity values of FWML and DAWM in SPMS participants^13^.

### 2.1. Population

Data used in this study is part of the International Progressive Multiple Sclerosis Alliance (IPMSA) study. We included MRI scans of 589 SPMS participants, enrolled in an unsuccessful clinical trial (no improvement of progression during the length of the trial), scanned longitudinally for 3 years, at screening, as well as weeks 24, 48, 72, 96, 108, and 156. The total number of timepoints included in this study was 3951^13^. All subjects had 3D T1w and 2D T2w images available. Table 1 summarizes the acquisition parameters for the T1w and T2w scans.

**Table 1.** MRI acquisition parameters. TR= Repetition time. TE= Echo time. TI= Inversion time. T1w= T1-weighted. T2w= T2-weighted. TE, TR, and Flip angle are expressed as ranges, to encompass all values used by different sites.

|  | **T1w** | **T2w** |
| --- | --- | --- |
| Sequence | 3D Flash | 2D TSE |
| Orientation | axial | axial |
| TR (ms) | 20-30 | 4800-6100 |
| TE (ms) | 3-11 | 78-89 |
| Flip Angle | 27-30 | 90-180 |
| Number of slices | 60 | 60 |
| Voxel size (mm) | 1×1×3 | 1×1×3 |

### 2.2. Image Processing

All MR images were processed using the following steps:

1. Image intensity non-uniformity correction using N3^14^.
2. Linear image intensity normalization into range [0-100].
3. Brain tissue extraction using brain extraction based on nonlocal segmentation technique (BEaST)^15^.
4. Tissue classification in the native space of the T1w images using a tissue segmentation pipeline based on a random forests classifier (BISON)^16^.
5. Rigid co-registration of the T2w images to the T1w images of the same timepoint. Using these transformations, the tissue masks and brain masks were also transformed to the native T2w space.
6. Linear registration (9 parameters) of the T1w images to MNI-ICBM152 template^17^.
7. Nonlinear registration of the T1w images to MNI-ICBM152 template^18^.
8. Generation of deformation-based morphometry (DBM) maps using MINC tools.
9. Using an atlas of the cortical and subcortical GM regions^19,20^, average Jacobian determinant values from the DBM deformation field were calculated as estimates of regional atrophy/change for each GM region.
10. Automatic segmentation of WM lesions using a Bayesian Classifier^21^.

All of the generated masks and registrations were visually assessed and those that did not pass this quality control (QC) were excluded from the analyses. All of the pipelines used in the processing of the images have been developed and validated for use in multi-center and multi-scanner datasets and have been previously used in many such applications^17, 16, 22^.

### 2.3. Manual Selection of Thresholds to Separate FWML and DAWM

Sixty-seven scans (timepoints) were selected from the dataset. To ensure the generalizability of the results, the subjects were selected to have MRIs from different scanner models, of three manufacturers (Achieva from Philips, Sonata, Symphony, and Avanto from SIEMENS, and SIGNA and DISCOVERY from GE). An expert rater (JM), reviewed the co-registered sequences for each timepoint and selected 2 T2w intensity threshold values to separate the NAWM from DAWM and FWMLs. These values were used to estimate the regression weights for the automatic thresholding technique for the entire dataset (see step 2.4 below).

### 2.4. Automatic Segmentation of FWML and DAWM

The main contrast used for separation of FWML and DAWM from NAWM tissue was the T2w image. To achieve this, the co-registered WM masks obtained from the tissue classification step were eroded by one voxel (in the axial plane) to avoid any partial volume effects between tissue borders (ventricular fluid and cortical GM). An initial NAWM mask was generated by excluding the WM lesions, generated with the Bayesian classifier, from this eroded WM mask. The mode of the intensity histogram of the T2w image within this initial NAWM was calculated. Previous work has shown that the manually selected thresholds and this automatically determined mode value have a linear relationship^12^. Using a linear regression, the optimal weights for estimating DAWM and FWML threshold values based on these automatically generated modes were calculated. Based on this criterion, any voxels within the eroded WM mask with intensities higher than 1.45 times the mode of the intensity histogram were labeled as FWML. Similarly, any voxels within the eroded WM mask in the range [1.15-1.45] times the mode of the intensity histogram were labeled as DAWM. The remaining tissue was labeled as NAWM. All of the separations were visually assessed by JM and those that did not pass this quality control were excluded (n=84 scans: 2%).

### 2.5. T1w-normalized Intensity Values in DAWM and FWML Areas

T1w-normalized values were calculated for the two areas of interest (DAWM and FWML) using the mode of the histogram of NAWM as the normalization factor. Due to the high correlation between MTR values and normalized T1w intensities, these two T1w-normalized intensities are adequate to reflect the degree of demyelination of a given area: the lower the intensity value, the greater the myelin loss.

### 2.6. Conversion of Lesion Types

Using the longitudinally registered first and last scans of each participant, FWML and DAWM masks of the two visits were compared, and voxels that changed from one mask type to the other between the two visits (e.g. from NAWM or DAWM to FWML) were identified (i.e. as NAWM-to-FWML, or DAWM-to-FWML). These labels were then propagated across all available timepoints and their T1w-normalized intensities were calculated.

### 2.7. Clinical variables

All participants were assessed for the following variables at the corresponding scan visit: 1) Global Expanded Disability Status Scale (EDSS) and its sub-scores (e.g. Cerebellar functional system), 2) Symbol Digit Modalities Test (SDMT), 3) 9-Hole Peg Test (9HPT), and 4) determination of confirmed disability progression (CDP) for 12 and 24 weeks. CDP was defined as an increase of the global EDSS score that remained stable for at least 12 or 24 weeks. The increase had to be of 1 point if the baseline EDSS was between 0 and 5.5, or 0.5 if the baseline EDSS was equal to or greater than 6.

### 2.6. Statistical Analysis

Demographic and MRI variables are presented using descriptive statistics, as either mean and standard deviation (SD), or median and range, according to their distribution. Differences in volumes and T1w normalized intensities of DAWM and FWML were assessed using related-sample Wilcoxon Signed Rank Test. Differences in volumes of DAWM-to-FWML, NAWM-to-FWML, and NAWM-to-DAWM between patients with CDP for 12 and 24 weeks (progressors) and patients without CDP (non-progressors) were assessed using Mann-Whitney U test.

Longitudinal mixed effects models were used to test the following hypotheses:

i. FWML and DAWM volumes change as a function of Disease Duration and Age at Onset.
ii. GM atrophy is associated with FWML and DAWM volume changes.
iii. Worsening clinical symptoms and progression are associated with FWML and DAWM volume changes.
iv. Longitudinal changes of mean T1w intensity values in DAWM-to-FWML voxels (DAWM areas on the first scan that become FWML on the last scan) are associated with progression.

DAWM and FWML volumes were log-transformed to achieve normal distributions. All continuous variables were z-scored prior to the analyses. Longitudinal mixed effects models were used to study the association between DAWM and FWML lesion volumes, regional GM atrophy measures, and clinical variables. All models included age at disease onset, disease duration, sex, and treatment (treatment did not have a significant effect in any of the analyses). *Subject* was entered as a categorical random variable. The regional GM-DBM results were corrected for multiple comparisons using false discovery rate (FDR)^23^ correction with a significance threshold of 0.05. All analyses were performed in MATLAB version R2017b.

## 3. RESULTS

From a total of 4035 scans, 84 were excluded due to misregistration of the T2w to the T1w image (2%). Figure 1 shows an example of the segmentation of DAWM and FWML for one subject. The first row shows axial slices of the T2w image. The second row shows the automatic tissue classification and WM lesion masks overlaid on the T2w image. The third row shows the automatically generated DAWM and FWML masks overlaid on the T1w images.

**Figure 1.**
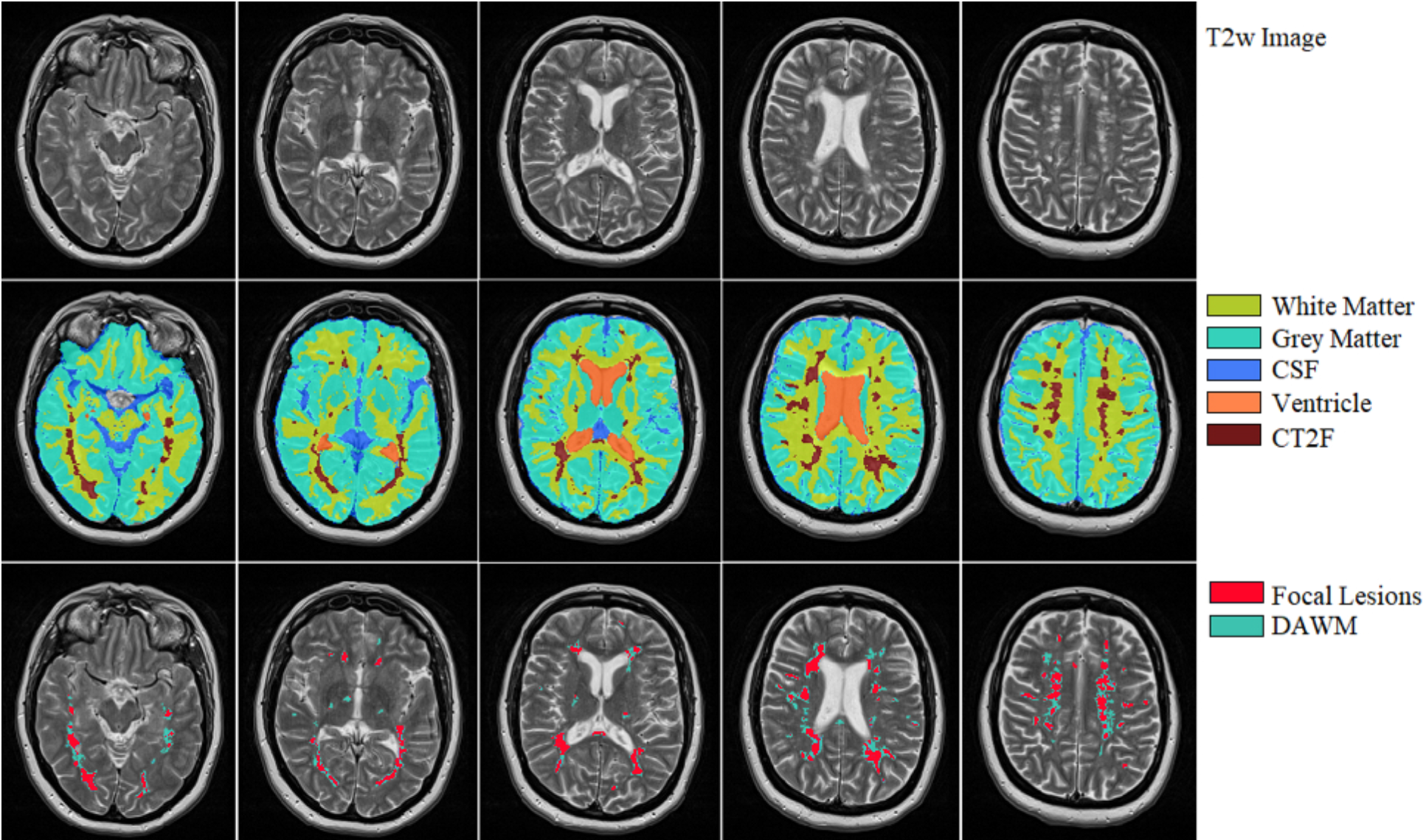
Example of the separation of DAWM and FWML. First row: T2w image. Second row: tissue classification and WM lesion masks on the T2w image. Third row: automatically generated DAWM and FWML masks on the T1w images. DAWM= Dirty Appearing White Matter. FWML= Focal White Matter Lesion. CSF: cerebrospinal fluid. CT2F: Bayesian lesion mask.

### 3.1. Population

The mean age of the participants at the time of the first scan was 49.24 ± 7.36 years, with a mean overall disease duration of 17.63 ± 7.32 years, and a duration of the secondary progressive phase of 6.38 ± 3.53 years. Table 2 summarizes the demographic characteristics of the participants.

**Table 2.** Demographic characteristics of the participants included in this study. Values represent number and percentage or mean ± standard deviation. RRMS= Relapsing Remitting Multiple Sclerosis. SPMS= Secondary Progressive Multiple Sclerosis. EDSS= Expanded Disability Status Scale.

|  |  |
| --- | --- |
| Sex | 245 M (36%) |
| Age at the time of scan | $49.24 \pm 7.36$ |
| Age at the onset of RRMS (age at first symptom) | $31.61 \pm 8.05$ |
| Age at the onset of SPMS | $42.86 \pm 7.34$ |
| Disease duration | $17.63 \pm 7.32$ |
| SPMS duration | $6.38 \pm 3.53$ |
| EDSS | $5.69 \pm 1.10$ |

### 3.2. MRI Variables

The median FWML volume in the dataset was 10.21cc (range: 0.03-13.61cc). The median DAWM volume was 21.59cc (range: 0.87-95.19cc) and it was significantly higher than the FWML volume (p<0.0001). The median FWML T1w-normalized intensity was 0.83 (range 0.00-1.00). The median DAWM T1w-normalized intensity was 0.96 (range: 0.01-1.01) and it was significantly higher than the FWML T1w-normalized intensity (p<0.0001).

### 3.3. Longitudinal FWML and DAWM Volume Changes

Mixed effects models were used to assess the relationship between FWML and DAWM volumes and Age of Onset and Disease Duration, controlling for Sex, and Treatment. Both FWML and DAWM volumes were significantly and negatively associated with Age at Onset: the older the person, the lower the volume of these two types of hyperintensity (p<0.0001 for both FWML and DAWM) (Figure 2, note use of ordinate log scale). FWML volume significantly increased (p=0.001) and DAWM volume significantly decreased (p<0.0001) as Disease Duration increased (Figure 2). Additionally, the ratio of FWML/DAWM volume per-participant significantly increased (p<0.0001) with Disease Duration (Figure 2).

**Figure 2:**
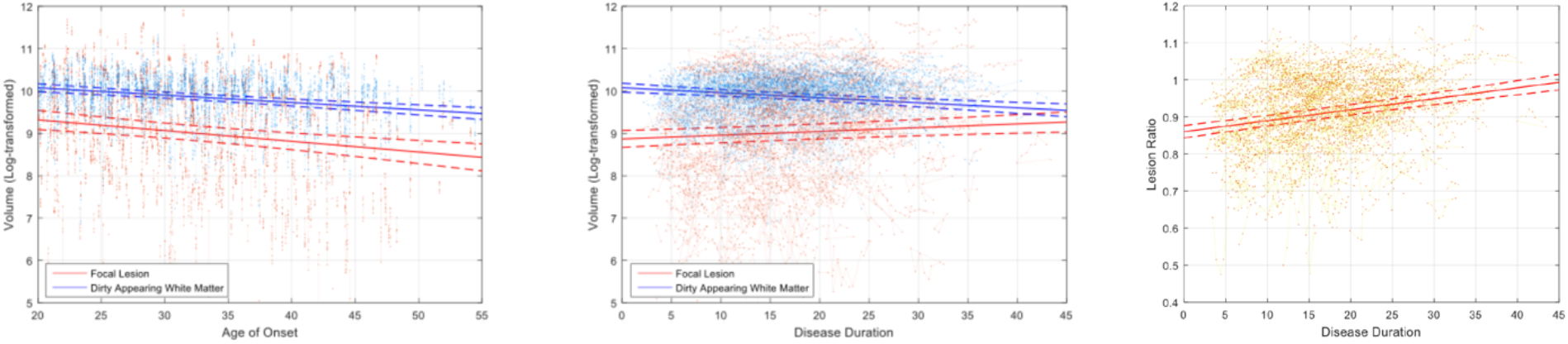
Left panel shows the volumes of DAWM (in blue) and FWML (in red) in relation to the age at disease onset (age at first symptom). Mid-panel shows the volumes of DAWM (in blue) and FWML (in red) in relation to the overall disease duration (from age at first symptom until the age at the time of the scan). Right panel: shows the Volume Ratio of FWML/DAWM in relation to disease duration

### 3.4. GM Atrophy and FWML and DAWM Volumes

Mixed effect models were used to assess the relationship between regional GM atrophy and FWML and DAWM volumes, controlling for Age, Sex, and Treatment. Figure 3 shows t-statistics maps of the GM regions that remained significant after FDR correction. Atrophy in both thalami, putamen, pallidum, brainstem, and cortical cerebellar and cerebral cortical GM were significantly associated with FWML volume increase (Figure 3). In parallel, cortical areas of right occipital lobe, right temporal lobe, left insular lobe, cerebellum, and brainstem were significantly associated with DAWM volume decrease (Figure 3).

**Figure 3:**
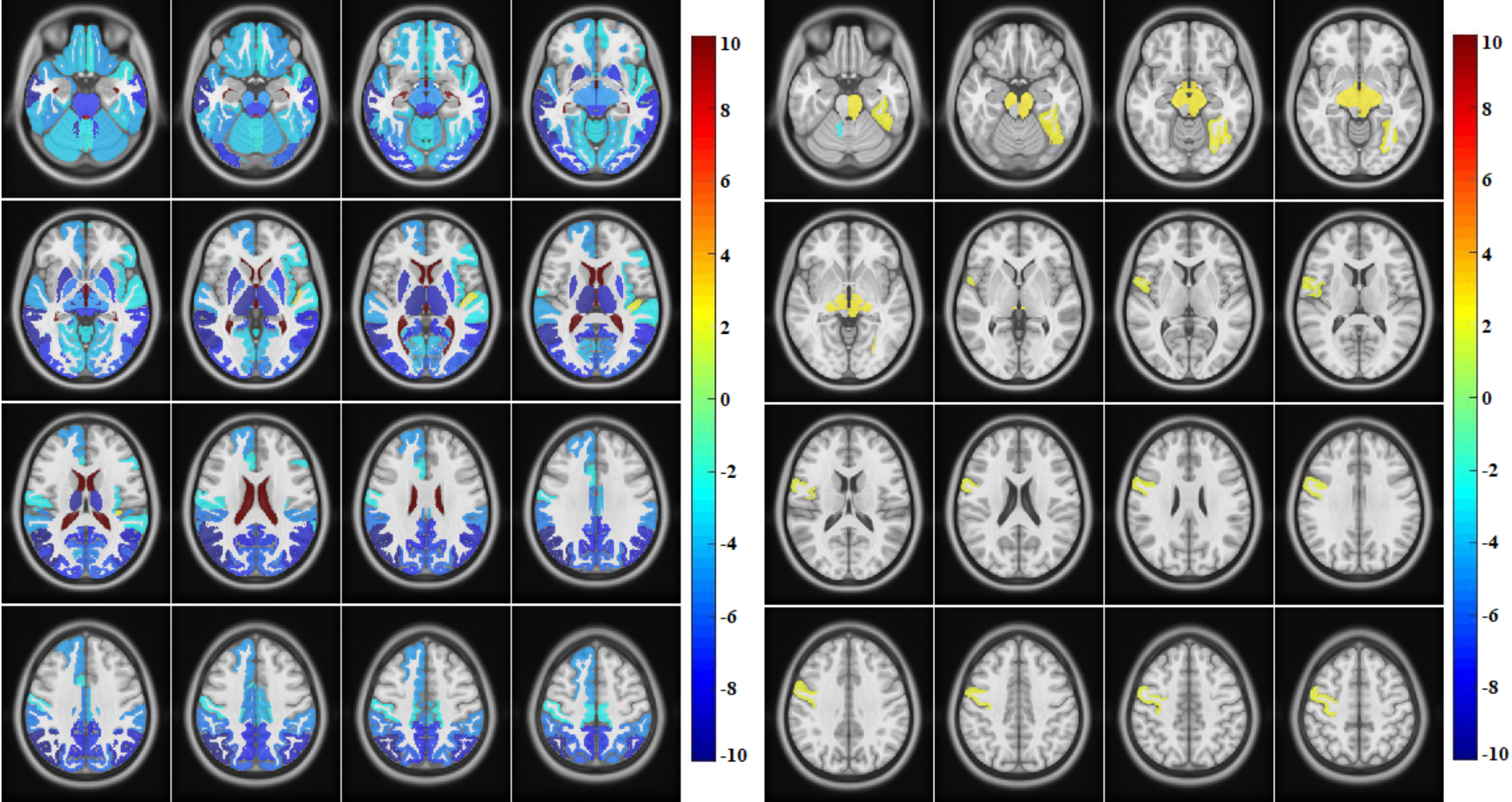
Left panel: t-statistical-map showing significant areas of atrophy associated to FWML volume increase. Right panel: t-statistical-map showing significant areas of atrophy associated to DAWM decrease.

### 3.5. EDSS, SDMT, and 9HPT scores as a function of FWML and DAWM volumes

Mixed effect models were used to assess the relationship between clinical variables and FWML and DAWM volumes, controlling for Age, Sex, and Treatment. Global EDSS scores were significantly and positively associated with FWML volumes (p=0.002), but not with DAWM volumes. Cerebellar, brainstem, and visual functional systems scores (EDSS sub-scores) were significantly and positively associated with FWML volume (p=0.002, p<0.0001, p<0.0001, respectively). Cerebral functional system score (EDSS sub-score) was significantly and positively associated with FWML volumes (p=0.002), and negatively with DAWM volumes (p=0.03). Other functional system scores (pyramidal, sensory, and bowel and bladder functional systems) showed no significant association with either FWML or DAWM volumes (Figure 4).

**Figure 4:**
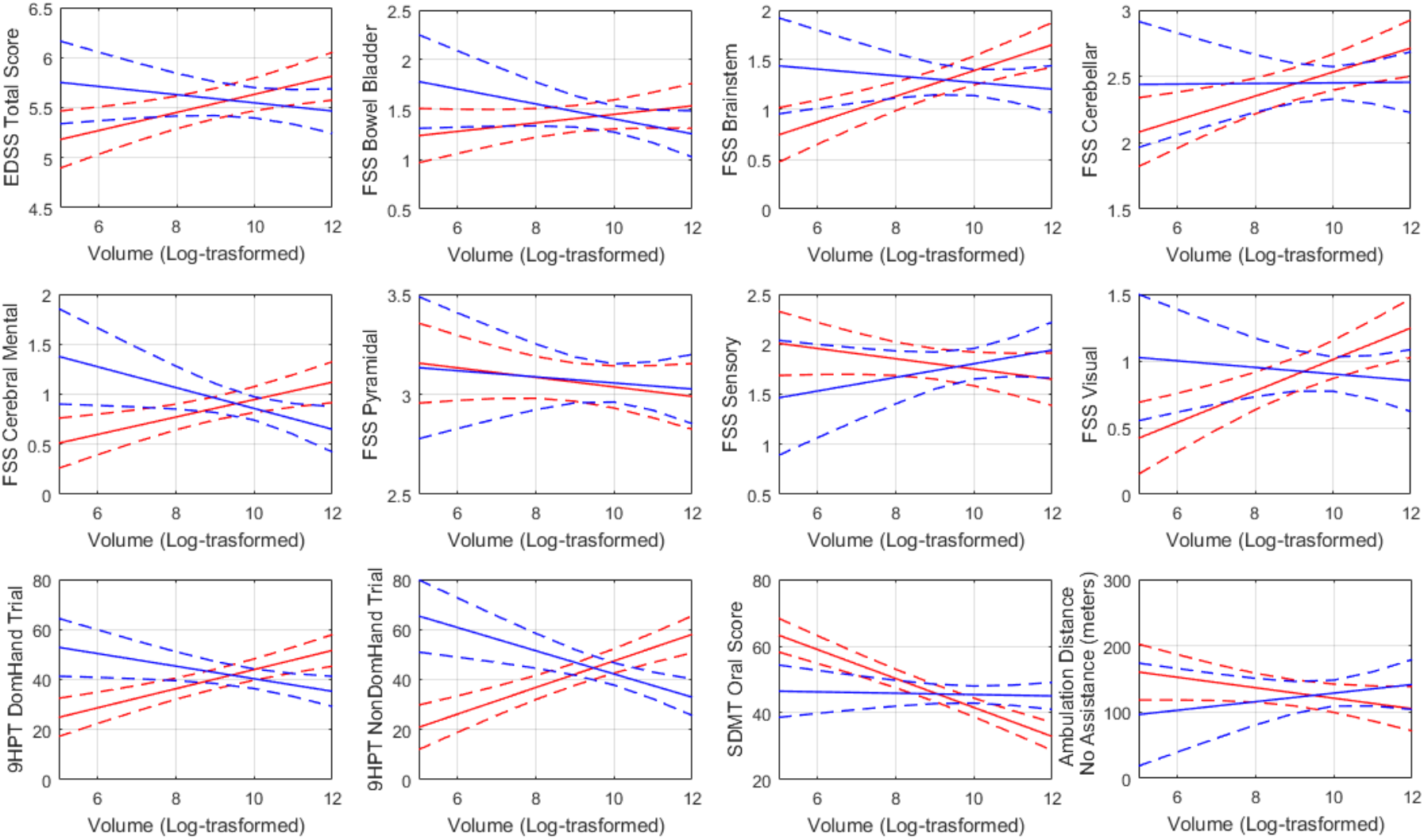
Association between DAWM and FWML volumes and clinical variables. Red lines: FWML volume. Blue lines: DAWM volumes.

The 9HPT time was significantly and positively associated with FWML, both in the dominant (p<0.0001) and non-dominant hands (p<0.0001). Significant and negative associations were also found for the dominant (p=0.02) and non-dominant (p=0.001) hand 9HPT results and the DAWM volumes. SDMT results showed a significant negative association only with FWML volumes (p<0.0001) (Figure 4).

### 3.6. FWML and DAWM volumes and Progression

CDP for 12 and 24 weeks showed a significant and positive association with increase of FWML volume during the course of the disease (p<0.0001 for both 12- and 24-weeks). DAWM volumes were not significantly associated with progression.

### 3.7. Evolution of DAWM-to-FWML and Progression

Voxels segmented as DAWM on the first scan of each participant that transformed to FWML on the last scan were considered as DAWM-to-FWML volumes and were significantly associated with progression (12-weeks p=0.01; 24-weeks p<0.001). Additionally, the median volume of tissue that transformed from DAWM-to-FWML was significantly higher in patients with CDP for 24 weeks (2.70cc vs 1.76cc; p<0.0001; F=3.1), representing 14% of the total DAWM volume at screening, compared to 10% in patients who did not progress. Similar results were obtained patients with CDP for 12 weeks (2.61cc vs 1.75cc; p<0.0001; F=1.5), representing 14% of the total DAWM volume at screening, compared to 10% in patients who did not progress.

Similarly, voxels of NAWM on the first scan of each participant that transformed to FWML on the last scan, NAWM-to-FWML volumes, were significantly higher in patients with CDP (12-weeks p=0.004; F=0.5; 24-weeks p=0.003; F=0.4). Also, voxels of NAWM on the first scan of each participant that transformed to DAWM on the last scan, i.e. NAWM-to-DAWM volumes, were significantly higher in patients with CDP at 24 weeks (p=0.01; F=0.06), but they did not reach significance for CDP at 12 weeks (p=0.06; F=0.001).

Conversely, voxels of DAWM that transformed into NAWM, or FWML that transformed into NAWM or DAWM did not show a significant association with progression.

Interestingly, visual inspection of the longitudinal changes in the DAWM and FWML regions showed that the areas of DAWM-to-FWML were mostly distributed in the perimeter of FWML, which is the reported location of activity of chronic active lesions^24^ (Figure 5).

**Figure 5:**
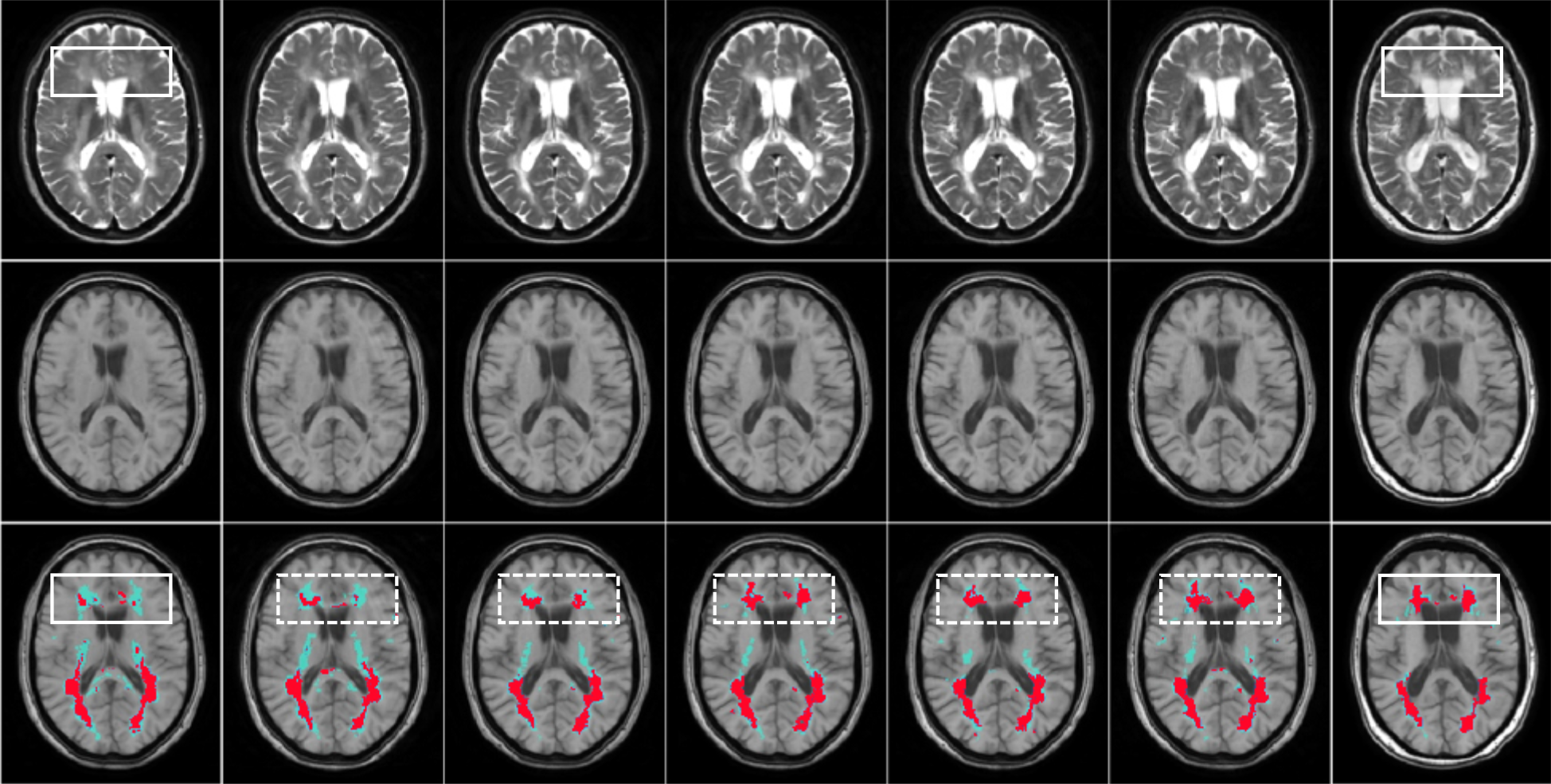
From left to right the columns represent the scans acquired at: screening, weeks 24, 48, 72, 108, 156. Top row: T2-weighted images. Mid row: T1-weighted images. Bottom row: T1-weighted with masks overlaying areas of FWML (cyan) and DAWM (red). Frontal periventricular FWML areas expand at the expense of DAWM located around the focal lesion (white box).

Finally, when assessing the evolution of T1w-normalized intensities in DAWM-to-FWML areas, and NAWM-to-FWML, both showed a significant negative association with 12- and 24-weeks progression (DAWM-to-FWML 12-weeks progression: p=0.004; DAWM-to-FWML 24-weeks progression: p<0.00001; NAWM-to-FWML 12-weeks progression: p<0.001, and NAWM-to-FWML 24-weeks progression: p<0.00001). T1w-noramalized intensities in NAWM-to-DAWM areas only showed a significant negative association with 24-weeks progression (p=0.006). Association of NAWM-to-DAWM with 12-weeks progression did not reach significance (p=0.07).

## 4. DISCUSSION

SPMS remains a form of MS that entails a poorer prognosis given the lack of response to most current available therapies that successfully control the focal inflammatory component of the disease^6, 12^. Different types of lesions, such as CLs ^25, 26^ and brain atrophy ^27, 28, 29^ have been proposed as being prominent in the progressive phase of MS, and hence considered as potentials predictors of fixed disability and progression. However, none have been identified as a definite predictive marker of progression. On one hand, CLs are difficult to identify on 3T MRI scans^30,31^, and require a highly experienced imaging expert to perform a very time-consuming manual segmentation. Additionally, 3T scans mainly capture a single type of CL, the leukocortical type, ^32,33^ and have very low sensitivity to detect subpial lesions^32^, the type that seems significantly associated with progression. In order to capture subpial lesions, 7T scans are necessary^32^, but such ultra-high-field MRI scanners are found in only a few centers and usually only used for research purposes. For these reasons, CL quantification seems impractical, at least for now, as a marker of MS progression. Brain atrophy has been another useful measurement, extensively explored as a marker of fixed disability^8^, but its exclusive role in progression is still difficult to weigh, given that atrophy rates seem to be similar in all stages of MS, from the clinical isolated syndrome phase, to the SPMS stage^34^.

Histopathology studies have suggested that the diffuse WM lesion component, DAWM, could be related to progression^7^. Our study explored not only the characteristics of DAWM, but also its relation to FWML at various levels, and proposes some of these characteristics as potential useful markers of progression:

- FWML areas showed significant lower T1w-normalized intensity values than those of DAWM areas, suggesting that DAWM exhibits a lower level of demyelination^12^.
- DAWM volume was significantly higher in patients that develop the disease at a younger age. In this respect DAWM did not differ from FWML (Figure 2). However, volume changes of FWML and DAWM were inversely related: as disease duration increased, DAWM decreased and FWML increased. We also calculated the ratio of FWML/DAWM volumes on a per-participant basis, in an attempt to assesses whether DAWM was turning into FWML on each individual case. We found a significant positive association of this ratio and disease duration.
- FWML volume increase and DAWM decrease were significantly associated to GM atrophy, with large areas of both, deep and cortical GM, related to FWML increase, and some areas of cortex and the brainstem related to DAWM decrease.
- FWML volume increase over time, but not DAWM decrease, was significantly associated to clinical variables such as EDSS and some of their sub-scores (Figure 4).
- FWML volume increase, but not DAWM decrease, was related to 12- and 24-week CDP.

Since the increase of FWML was more prominently associated with clinical variables, but, at the same time, the FWML/DAWM ratio suggested a change of DAWM into FWML volumes on a per-participant basis, we explored associations of voxels that were segmented as DAWM in the first scan and transformed to FWML in the last scan. We then related this volume (i.e. DAWM-to-FWML volume) with progression. We found that DAWM-to-FWML volume was significantly higher in patients with CDP (for both 12- and 24-weeks). NAWM-to-FWML volume was also significantly higher in CDP (for both 12- and 24-weeks), although with lower F values than DAWM-to-FWML. NAWM-to-DAWM showed a significant difference only in participants with CDP for 24weeks, and was marginally significance for CDP for 12 weeks (p=0.06). These results suggest that conversion of DAWM to FWML seem to seal the progressive course.

We also noted that the distribution of the DAWM-to-FWML voxels occurs in the perimeter of FWMLs, suggesting a potential overlap with areas that have been described as chronic active lesions by other groups^24^.

Finally, we explored the changes in T1w-normalized intensity values in the areas of DAWM-to-FWML and NAWM-to-FWML. They both showed significant negative associations with 12- and 24-week CDP. Moreover, NAWM-to-DAWM T1w-normalized intensity values, also significantly decrease, showing a negative association with 24-week progression. The progressive decrease of T1w-normalized values as the disease progresses and their significant association with progression highlights the impact of the severity of these lesions (e.g. stronger demyelination). To summarize, not only a significant quantitative volume change of DAWM-to-FWML and NAWM-to-FWML characterized patients who progress, but also a significant decrease in T1w-normlized values of these areas, which translates the qualitative aspect of the change: more severe demyelination in these areas.

Our study is not without limitations. There were no quantitative MRI sequences such as magnetization transfer ratio (MTR) images or diffusion available, to assess the nature of the changes in a given area in a more quantitative manner. However, previous studies have shown a significant association between MTR and T1w-normalized values^12^. Another limitation is the lack of scans of these participants during the relapsing-remitting course of the disease. We acknowledge the possibility of different types of changes for DAWM and/or FWML, earlier in the course of the disease.

## CONCLUSION

FWML volume was significantly associated with GM brain atrophy, 9HPT, SDMT, specific EDSS functional systems (e.g. cerebellar), and progression in this group of SPMS participants.

DAWM transformed into FWML over time, and this transformation was significantly associated with clinical progression. DAWM voxels that transformed had greater normalized T1w intensity decrease over time, in keeping with relatively greater tissue damage evolution. Therefore, evaluation of DAWM in progressive MS may provide a useful measure for therapies that aim to protect this at-risk tissue, which would have the potential to slow progression.

